# In silico analysis of the human titin protein (Immunoglobulin-like, fibronectin type III, and Protein kinase domains) as a potential forensic marker for postmortem interval (PMI) estimation

**DOI:** 10.64898/2026.03.06.710245

**Authors:** Muhammad Umer Gill, Maleeha Akhtar

## Abstract

Due to the limited availability of reliable and well-validated molecular markers, the determination of postmortem interval (PMI) is still a major obstacle for forensic investigators to resolve a case. The largest human protein, known as titin, has never undergone at domain level examination of postmortem degradation patterns. This study focused on the In-silico analysis of the Immunoglobulin-like, fibronectin-type III, and Protein kinase domains of human titin to assess their potential utility in PMI estimation. Sequence data for the studied domains were retrieved from UniProt, 2D & 3D models were generated by PSIPRED and SWISS-MODEL, respectively, followed by physicochemical properties, solubility assessment, and structural comparison. This study revealed that the Ig-like domain is the most stable, followed by the Fn-III and Protein kinase domains. These findings indicate that Titin domains may degrade at different rates in the postmortem period. This study introduces the first computational basis for considering Titin as a multi-domain candidate biomarker for PMI estimation, laying the groundwork for upcoming laboratory validation.

## 1. Introduction

Determining the time of death is one of the most common and curioustic questions of crime scene and forensic laboratories. Authentication and revocation of testimony in crimes, factors of death, and ultimately for the solution of crimes a reliable estimation of the time since death is essential [1]. Primary changes after death include the development and remission of Rigor mortis, advancement of livor mortis, and algor mortis [2]. Alternative strategies to analyze changes in early postmortem Interval (PMI) encompass ocular reactivity to pharmacological stimulations and electrostimulation of specific muscles [3,4] are the commonly used methods to demarcate PMI in forensic work, but there are many errors and constraints in most of the cases. Vitreous potassium and similar biochemical markers exhibit considerable variability influenced by environmental and biological factors, including temperature, physiology, and agonal state[5]. Similarly, postmortem protein degradation, show effective in experimental studies, but remain poorly standardized and limited validation for practical forensic applications in humans[6]. Therefore, proteomic and computational analysis are used to allow detailed analysis of protein fragments, protein turnover and degradation providing forensic investigators with advanced molecular markers that surpass the limitations of classical morphology and biochemical techniques [7]. Analyzing the postmortem degradation of proteins in skeletal muscles has emerged as promising approach to predict the time since death, particularly with the early to mid-postmortem interval (24-124 hours). However within these proteins base methods only few proteins (Titin, Desmin, Nebulin and Tropomyosin) are used to study proteolysis after death[7]. This research mainly focuses on the domains of the Titin protein such as Immunoglobulin-like (Ig-like Domain), Fibronectin type III (Fn-III Domain) and Protein kinase domain, because degradation rate analysis of certain proteins, including titin, appears to be reasonably consistent, serving as a basis for protein-specific or domain-specific molecular chronometers for PMI [8]. The main reason of choosing these specific domains of the titin protein includes the different degradation time of each domain and different processes of degradation by different cellular events [9].To the best of current studies this is the first computational analysis focusing on the Ig-like, Fn- III, and Protein kinase domain of titin may serve as a promising molecular markers for the assessment of PMI.

## 2. Materials and Methods

This study utilizes a comprehensive bioinformatics framework involving sequence collection, 3D-structural modeling, and physicochemical profiling of the studied domains. We have performed all procedures using standardized protocols and widely accepted computational tools.

### Sequence Retrieval

Sequences corresponding to the human titin, Immunoglobulin-like, Fibronectin type III, and Protein kinase domains were obtained from the UniProt database (Q8WZ42) (reference genome). The retrieved sequence in FASTA format served as the basic dataset for all subsequent analyses.

### Selection Criteria and Boundary Determination

Human titin protein is composed of repeated immunoglobulin-like (Ig) and fibronectin type III domains which together constitute the majority of protein sequence and play an important role in sarcomere architecture and elasticity [10]. Individual Ig and Fn-III domains exhibit conserved folding architectures, enabling domain-level analysis despite extensive sequence repetition. For example, consensus structural alignment based on experimentally determined structure, along with multiple high-resolution Fn-III domain entries in the Protein Data Bank (PDB), support their use as validate representative template. Therefore, to achieve computational tractability without compromising biological significance, a single representative Ig-like (6-96) and one FN-III (14019-14114) were selected based on curated structural annotations and positional representation within titin molecule. The unique protein kinase domain (residue 32178-32432) was analyzed in it’s entirely, given its singular occurrence. The adopted domain selection strategy allows comparative evaluation of domain specific stability and degradation propensity across domains, while remaining consistent with established structural knowledge of titin.

### Physicochemical Characterization

ExPASy ProtParam tool was used to calculate the physicochemical properties [11] of the studied domains that include number of amino acids, their molecular weights (kDa), theoretical pI at which the net charge is zero, instability index, aliphatic index, GRAVY scores and estimated half-life. The given parameters are used to analyze the stability of protein domains, their hydrophobic characteristics, and the way by which each domain behaves under physiological conditions.

### Predicted Solvent Accessibility & Secondary Structure Features

Amino acids exposure to surface and the percentage of amino acids that are buried inside is termed as solvent accessibility and it is calculated by using I-TASSER tool [12] which is used to predict protein structure and functions. The resultant scores ranges from 0 to 9 representing highly buried and highly exposed residues respectively. However, well know tool PSIPRED was used to predict the secondary structure content of the studied domains showcasing α-helix, β-sheets, and loop/coils.

### 3D Structure Predictions

SWISS-MODEL was used to predict the 3D structure [13] of the studied domains. There were many structures; the most reliable structure was selected based on the global model quality estimation (GMQE) and qualitative model energy analysis (QMEAN). PDB files were exported for the most reliable and accurate structures and used for further structural evaluation.

### Structural Validation

The structural stability of each exported model was verified through these given tools.

1. The **Ramachandran Plot** generated by Swiss-Pdb viewer was used to study stereochemical quality.
2. The **ERRAT2** was used to analyze non-bonding interaction quality.

### Hydrogen Bond Analysis

Chimera version 1.19 was used to study the domain PDB files to visualize and study the hydrogen bond. Donor-acceptor distance and angles were used to map the hydrogen bond in each domain. This analysis provides insight into the structural integrity and stability of the titin domains.

**Figure 1.**
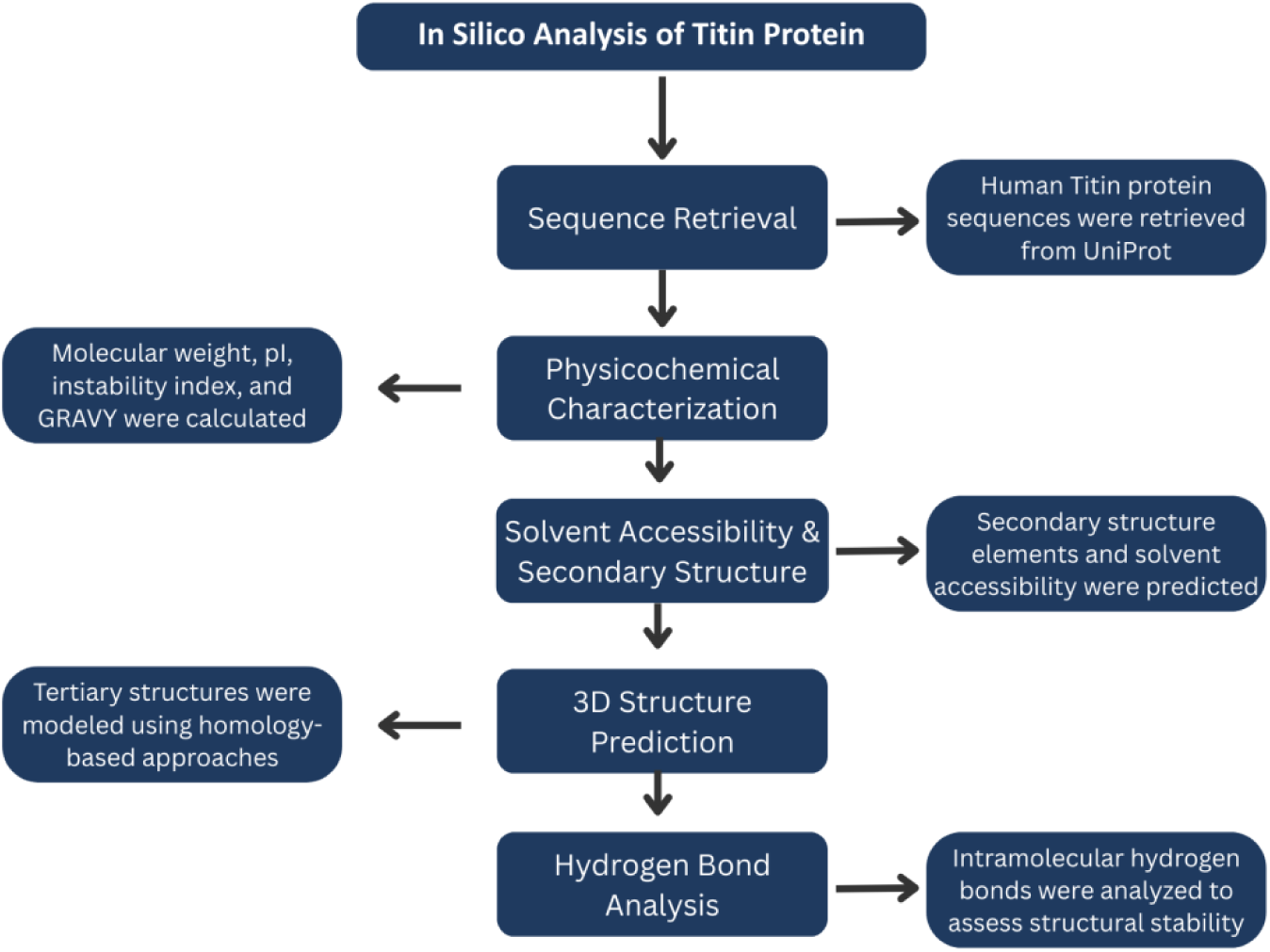
Workflow of the in silico analysis performed for the human Titin protein. The flowchart illustrates the sequential steps including sequence retrieval from UniProt, physicochemical characterization, solvent accessibility and secondary structure prediction, three-dimensional structure modeling, and intramolecular hydrogen bond analysis to evaluate structural stability.

## Results and Discussion

### Physicochemical Characterization of titin protein domain

Physicochemical properties, including the number of amino acids, molecular weight (kDa), theoretical isoelectric point (pI), instability index, aliphatic index, GRAVY score, and estimated half-life in mammalian cells, are discussed in..

**Table 1.**
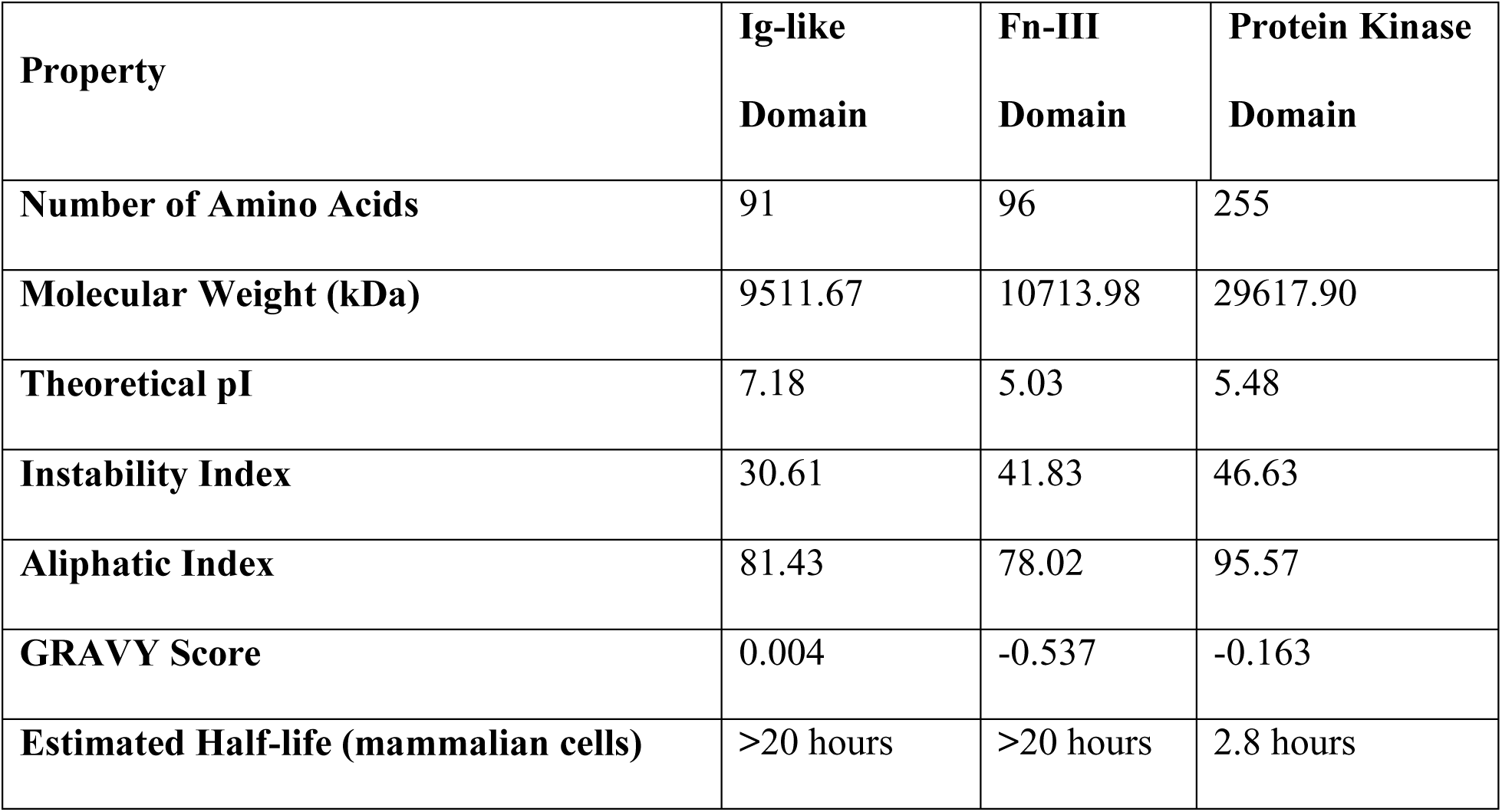
Comparative physicochemical profiling of Ig-like, FN-III, and Protein kinase domains of the human titin protein. Key parameters relevant to domain stability, hydrophobicity, and degradation susceptibility, such as molecular weight, theoretical pI, instability index, aliphatic index, and GRAVY score, are shown in the this table. The ExPASy ProtParam server computed all listed physicochemical parameters.

The number of amino acids and corresponding molecular weight reflect the primary structural profile of each domain, and higher-molecular-weight proteins typically undergo more complex degradation patterns after death. The theoretical pI determines the protein’s net charge at physiological pH, influencing its solubility, its tendency to aggregate and overall susceptibility to proteolytic degradation after death. Proteins possessing extreme pI values may exhibit increased precipitation or show altered degradation kinetics under postmortem conditions [14]. The instability index reflects a protein’s propensity for degradation in vitro, earlier research reported that proteins exhibiting higher instability indices undergo accelerated degradation during the postmortem interval [15]. The aliphatic index reflects the relative content of aliphatic side chains, correlates with the thermal stability of the proteins. Proteins exhibiting higher aliphatic indices enable proteins to maintain their structure in challenging environments, increasing their postmortem persistence in forensic investigations[16]. The GRAVY (Grand Average of Hydropathy) score indicates a protein’s overall hydropathicity, that hydropathic proteins are generally resistant to degradation by enzymes in an aqueous environment yet remain susceptible to aggregation [17]. Finally, the estimated half-life in mammalian cells provides an estimation of protein longevity, but in reality, postmortem stability is affected by protein physicochemical properties and environmental factors [18,19]. On the basis of the above-discussed parameters, the instability index and aliphatic index are the most important factors, thus it can be concluded that the protein kinase domain is the most structurally stable, followed by the Fn-III domain and the Ig-like domain.

### Solvent accessibility

Analysis of solvent accessibility estimates distinct variations in the proportion of buried and surface–exposed residues among the three titin domains [20]. Studies indicate that the sequestration of non-polar solvent-accessible residue is a major factor governing the protein stability, underscoring how hydrophobic burial stabilizes and strengthens the core protein formation. With this background stability difference, the domain-wise results are described in Figure 2.

**Figure 2.**
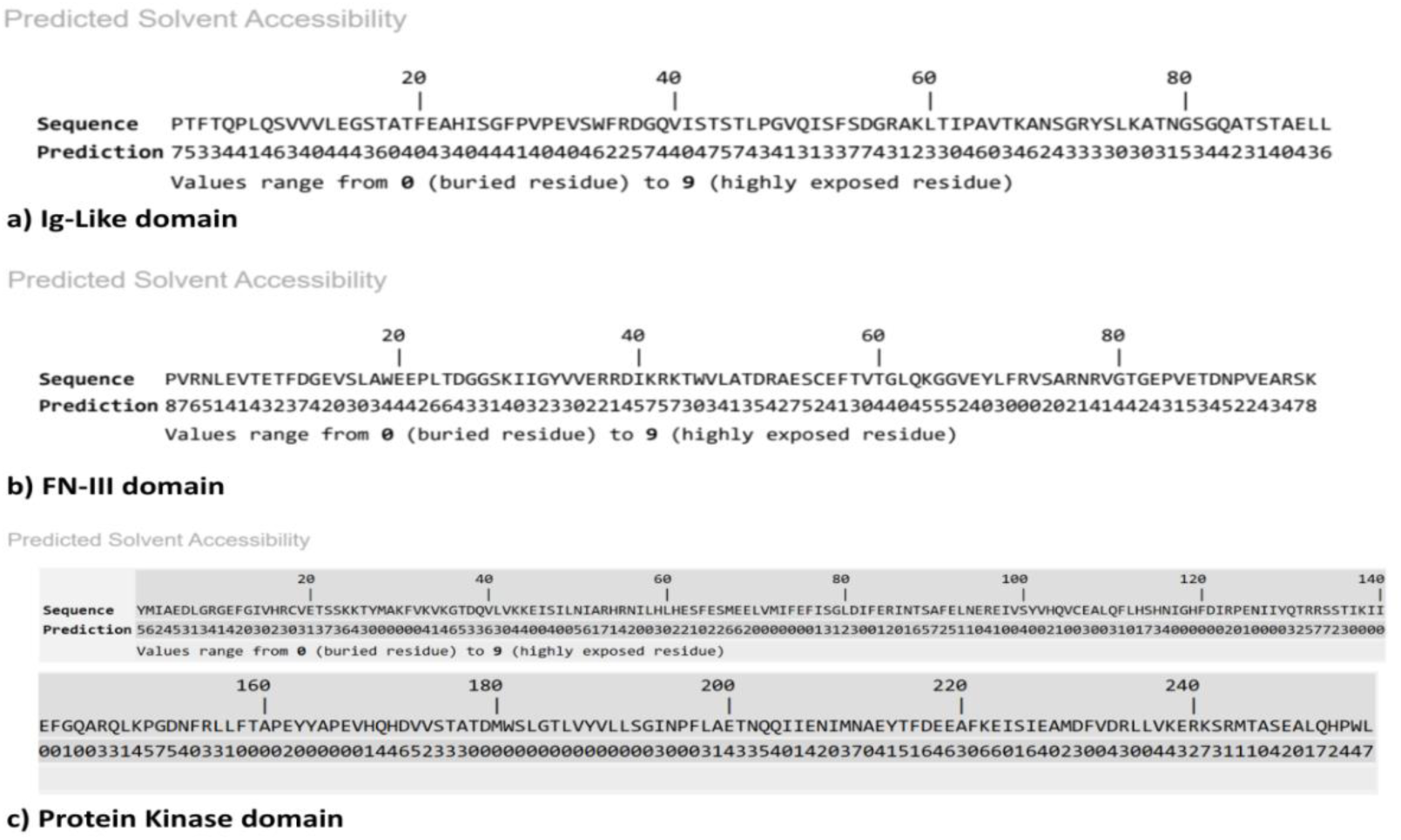
The predicted solvent accessibility of selected domains by using the I-TASSER model. Sequence a) shows the predicted solvent accessibility of Ig-like domain, b) represents Fn-III domain, and c) represents the Protein kinase domain. Residue exposure levels were predicted by using I-TASSER structural model. This profile reveals accessible loop areas, whereas the β-sheet core is predominantly buried, reflecting the compact architecture of this domain.

Analysis of solvent-accessibility pattern in **Error! Reference source not found.** shows that Ig-like domain contains an appropriate distribution of buried (0-2) and surface-exposed (6-9) residues, indicating the typical organization of a β-sandwich fold. Low-accessibility residues indicate a compact hydrophobic core that reinforces structural stability, while highly exposed residues are mainly located in surface loops. Similarly, for Fn-III domain which exhibit a combination of buried and exposed residues with several segments exhibiting high solvent accessibility scores (≥7), mainly found in loop or terminal regions. Buried residues (0-2) form a stable core, while exposed regions may be more susceptible to postmortem degradation. Lastly, the protein kinase domain also contains an appropriate amount of buried residues, which contains most of the values ranging from 0-3, representing the buried residues and some extent of residues having values ranging from 4-6, and a very low amount of residues with values greater than 7.

### Secondary structure features

The secondary structure features of the selected domains that describe the local arrangement of the protein backbone and are classified into α-Helix, β-Strand, and coil/loop regions. These structural elements are governed by the backbone hydrogen bonding, which reinforces the stability of the protein fold. α-Helix and β-Strands serve as structural motifs that provide order and stability, in contrast to loop and coils, which constitute flexible, dynamic and adaptable regions that are less stable yet functionally significant [21,22]. In this study, secondary structure predictions were obtained using PSIPRED, a high-accuracy, neural network- based tool commonly used to classify each residue into helix, strand, and coil according to sequence-based features [21]. The structural analyses of Ig-like, fibronectin Type III, and protein kinase domains are illustrated in Figure 3, Figure 4, Figure 5 respectively.

**Figure 3.**
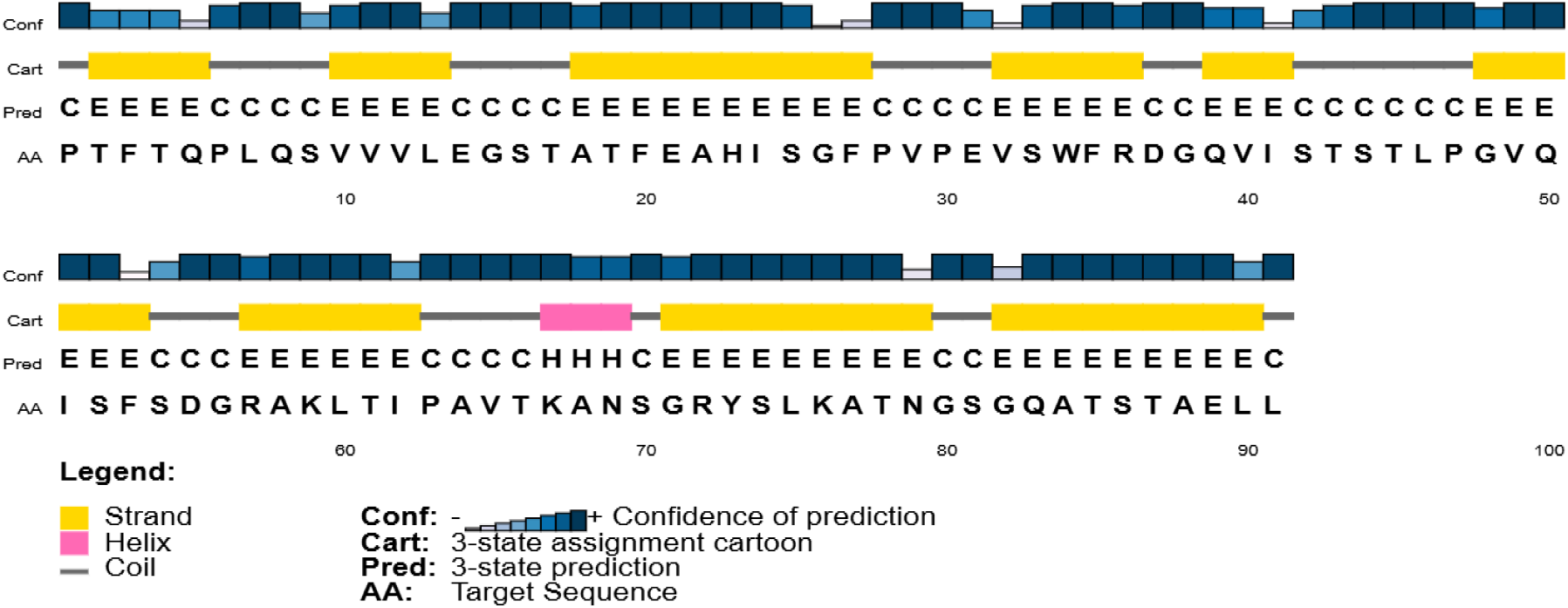
Illustration of predicted secondary structure of immunoglobulin-like domain of human titin protein which was generated by PSIPRED tool. This Figure 3 shows the arrangement of the β-strands (Yellow), α-helix (Pink), and coil/loop regions (grey) beside their linear sequence. This distribution highlights the existence of high numbers of β-sheets, which provide the structural stability.

**Figure 4.**
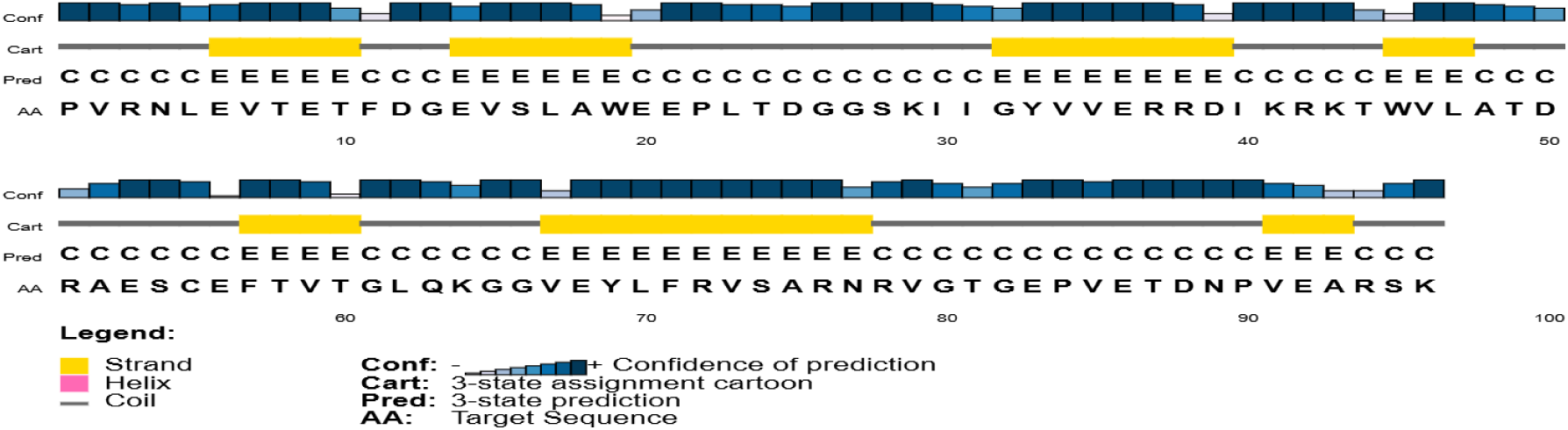
Illustration of Predicted secondary structure of Fibronectin Type III domain of Human Titin protein which is formed by PSIPRED tool. This Figure 4 shows the arrangement of the β-strands (Yellow), α-helix (Pink), and coil/loop regions (grey) beside their linear sequence. This distribution highlights the existence of high numbers of β-sheets, which provide the structural stability.

**Figure 5.**
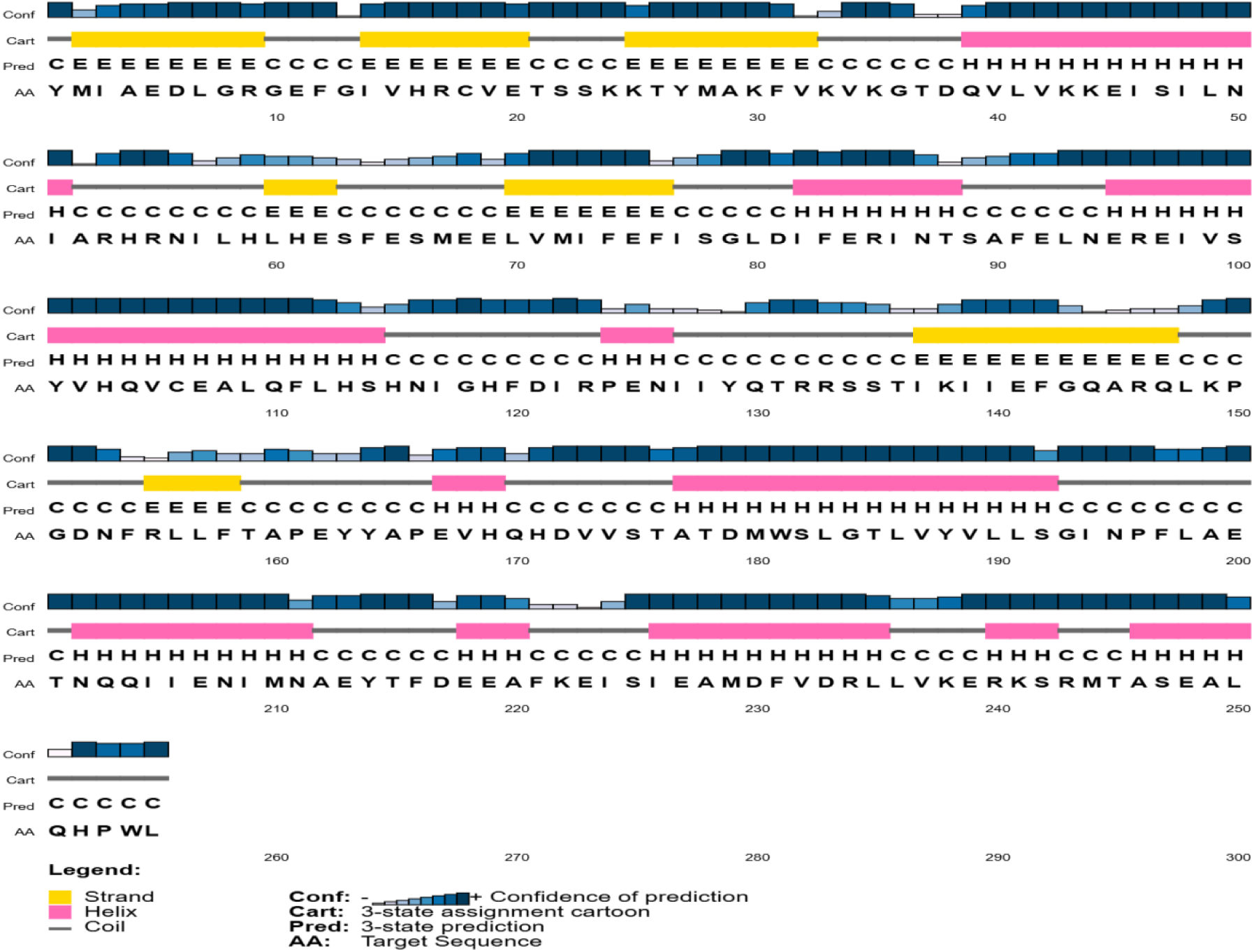
Illustration of Predicted secondary structure of Protein Kinase domain of Human Titin protein which is formed by PSIPRED tool. This Figure 5 shows the arrangement of the β-strands (Yellow), α-helix (Pink), and coil/loop regions (grey) beside their linear sequence. This distribution highlights the existence of high numbers of β-sheets, which provide the structural stability.

The secondary structure profile relies on the classification of residues in α-helices (pink), β-strands (yellow), and coils (grey). Regions adopting α-helices and β-strands tend to demonstrate greater structural stability while coil regions exhibit greater flexibility and tend to degrade earlier. Therefore the order of stability among these domains was evaluated on this basis β-strands > α-helices > coils indicating that fibronectin type III degrade rapidly followed by protein kinase and Ig-like domains respectively[23].

### 3D-Structure

The 3D structure of protein describes the precise spatial organization of amino acid residues, resulting in defined folding elements such as α-helices, β-sheets, and multi-domain architecture [24]. This multi-level structural organization is essential for governing protein stability, functional performance, and vulnerability to degradation, as proper biological activity depends on its native conformation [25]. Alteration of the native 3D conformation, whether from mutation, denaturing conditions, or postmortem effects, leads to functional instability and protein destabilization, making it more prone to degrade [24]. The subsequent images in Figure 6 demonstrate the 3D conformation and effects of such alterations.

**Figure 6.**
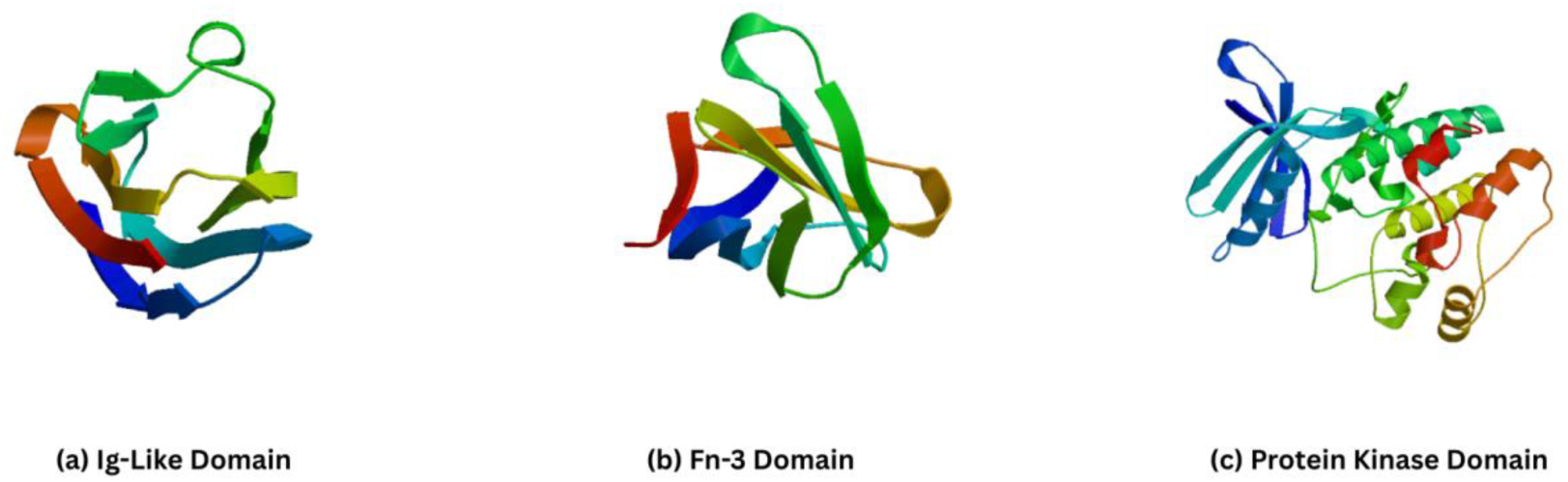
Three-Dimensional structure of Ig-like, Fn-III and Protein kinase domains of human titin protein. (a) Ig-like domain, (b) Fibronectin Type III domain (c) Protein Kinase domain respectively. Each domain is shown in a rainbow color scheme from N-terminus region (blue) to C-terminus (red)

In these 3D structures (Figure 6), the arrows indicate β-sheets oriented from the N-terminus to the C-terminus of the strand, helping to visualize the orientation within the sheet. Similarly, the lines joining the arrows represent the Loops/Coils, where the backbone is more flexible and does not form regular secondary structure [26]. Each 3D model highlights the distribution of β-sheet framework and loop/coil regions, influencing the domain stability and flexibility, respectively. These structural features correspond with physicochemical data listed in **Error! Reference source not found.**, from which the comparative stability order is derived.

**Table 2.** Represents the relative stability of Immunoglobulin-like, fibronectin type III, and protein kinase domain of human titin protein; based on structural features, physicochemical properties, and three-dimensional structure. Comparative structural features, physicochemical properties, and stability assessment of the Immunoglobulin-like (Ig-like), Fibronectin type III (FnIII), and Protein Kinase domains of human titin. The table summarizes β-sheet and loop/coil composition, instability index, aliphatic index, GRAVY score, and predicted half-life to evaluate relative structural stability and flexibility among the three domains.

| Domain | Structural Feature<br>(β-sheets & loop/coils) | Key Physicochemical<br>Properties | Stability Assessment |
| --- | --- | --- | --- |
| Ig-like | • Very high β-sheet content.<br>• Minimum coils/loops | • Instability index: 30.61(Lowest)<br>• Aliphatic index:81.43<br>• Gravy: 0.004<br>• Half-life > 20h | Highly stable, compact β-sheet framework supported by favorable physicochemical properties. |
| Fibronectin type III | • High β-sheet content<br>• Few loops, mainly rigid | • Instability index: 41.83(Moderate)<br>• Aliphatic index:78.02<br>• Gravy: -0.537<br>• Half-life > 20h | Stable, dense β-sheet packing, least flexible regions, and supported by physicochemical properties as well. |
| Protein Kinase | • Moderate β-sheet content<br>• Many loops/coils, high flexibility | • Instability index: 46.63(Highest)<br>• Aliphatic index: 95.57(Highest)<br>• Gravy: -0.163<br>• Half-life: 2.8 h | Least stable, much more loops, minimal half-life, and highest aliphatic index. |

### Structural Validation and Analysis

The structural stability of the predicted domain models was performed through detailed 2D and 3D analyses to ensure the accuracy and reliability of these structures.

### Ramachandran Plot

The assessment of 3D models’ stereochemical quality and structural feasibility through a Ramachandran plot was done. The Ramachandran plot visualizes backbone dihedral angles (ψ and φ), verifying that residues fall within energetically favored regions. It enables us to identify localized geometric deviations and ensure the structural accuracy and stereochemical quality of protein modeling prior to downstream analyses. This method is widely regarded as a standard practice for confirming protein geometry [27,28]. Ramachandran plots for the studied domains are given in Figure 7.

**Figure 7.**
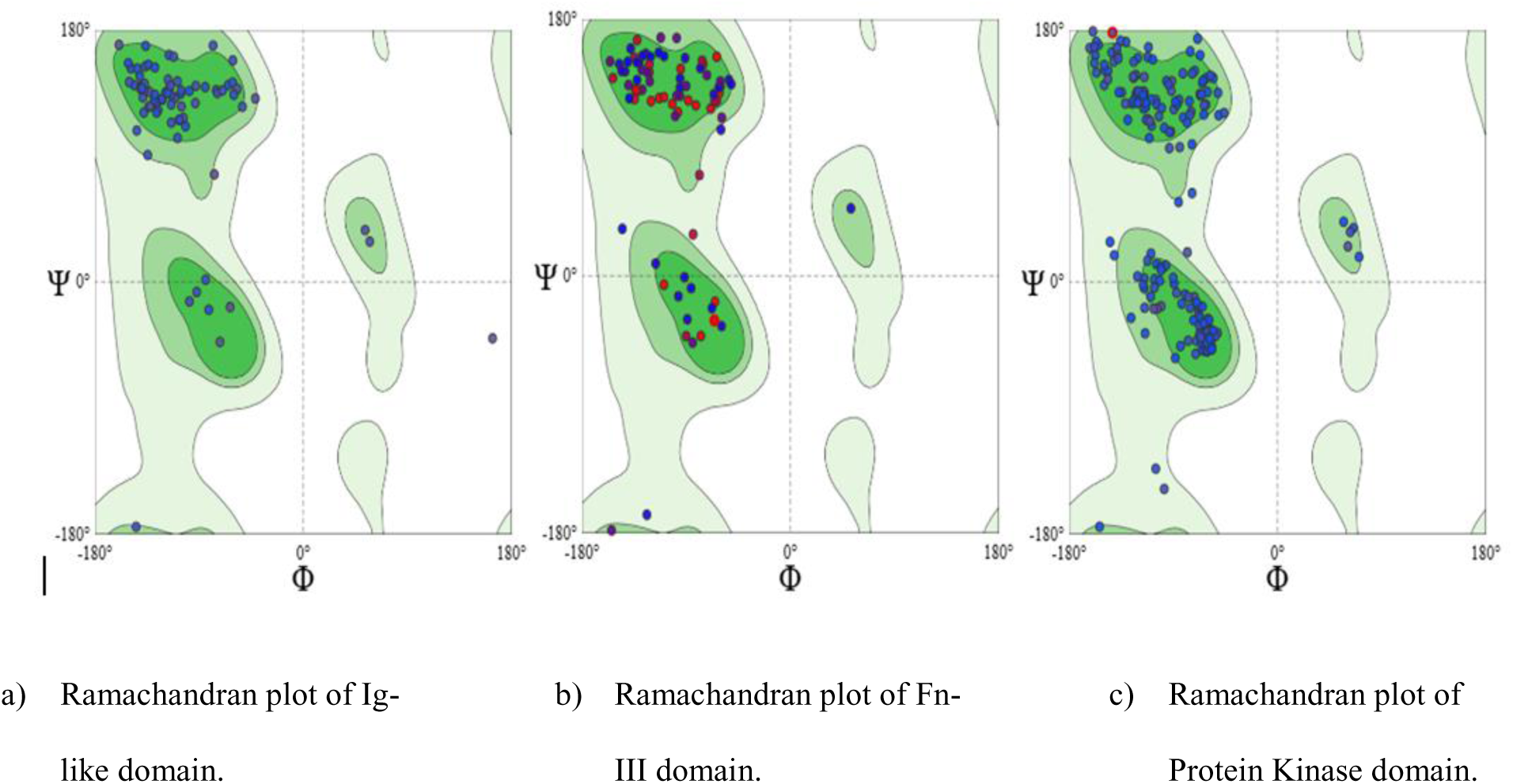
Structural validation of the titin protein model using Ramachandran plot analysis and quality assessment parameters. (a) Ig-like domain, (b) Fibronectin Type III domain (c) Protein Kinase domain respectively. These graphs were predicted by using SWISS-MODEL (ExPASy) server.

The dots in these graphs in Figure 7 represent the individual amino acid residues of each domain; the x-axis and y-axis represent the (ϕ, phi) rotating angle around the N-Cα bond and the (ψ, psi) rotating angle around the Cα-C bond, respectively. The Table 3 clarifies the favored, allowed, and outlier regions fall on a Ramachandran Plot [29].

**Table 3.** Summary of Ramachandran plot regions; illustrating the characteristic ϕ and ψ backbone dihedral angel ranges for favored, allowed and outlier regions. These regions represent the stereochemical validity and their relationship to typical secondary structural motifs. Although the Ramachandran plot does not directly determine the stability of human titin protein domains, more residues in the favored regions indicate both energetic stability and minimal geometric strain. So, according to Figure 7 Ig-like domain contain almost all the residues in the favored regions, while Fn-III domain has some of its residues in allowed regions and protein kinase domain has much of its residues in allowed regions.

| Region | Description | Approximate<br>phi($\phi$) angles($^{\circ}$) | Approximate<br>psi($\psi$) angles($^{\circ}$) | Typical<br>Secondary<br>Structure |
| --- | --- | --- | --- | --- |
| **Favored** | Most stable and frequently observed | a) $-60^{\circ}$ to $-120^{\circ}$(α- | a) $-30^{\circ}$ to $-60^{\circ}$(α-Helix) | Represents α-Helix, β-sheet |
| | backbone conformation. | Helix),<br>b) $-120^\circ$ to $-180^\circ$(β-Sheet) | b) $120^\circ$-$180^\circ$(β-Sheet) | and left-handed helix. |
| **Allowed** | Less optional yet structurally acceptable region. | a) $180^\circ$ to $-60^\circ$ ($\alpha/\beta$ transition) | a) $-180$ to $180$ (varied and less common) | Loops, turns and less common regions. |
| **Outlier** | Outlying residues reflecting energetically unfavorable geometry. | a) Any $\psi$ or $\phi$ outside the allowed or favored region fall in this category. | 1. Any $\psi$ or $\phi$ outside the allowed or favored region fall in this category. | Modeling errors or unusual conformations. |

### ERRAT 2

Our selected domains structures were validated by using ERRAT 2, a structural-evaluating tool from institute of genomics and proteomics under the U.S. Department of Energy Office of Sciences. This tool evaluates the structure based on analysis of non-bonded atomic interactions. The results appeared in form of percentage that indicated the reliability of the structure, with higher or better percentage indicating minimal stereochemical errors[30]. This analysis served as a supplement to Ramachandran analysis by revealing additional localized structural deviations. The graphical representation of each domain’s ERRAT 2 analyses is given in **Figure 8**.

**Figure 8.**
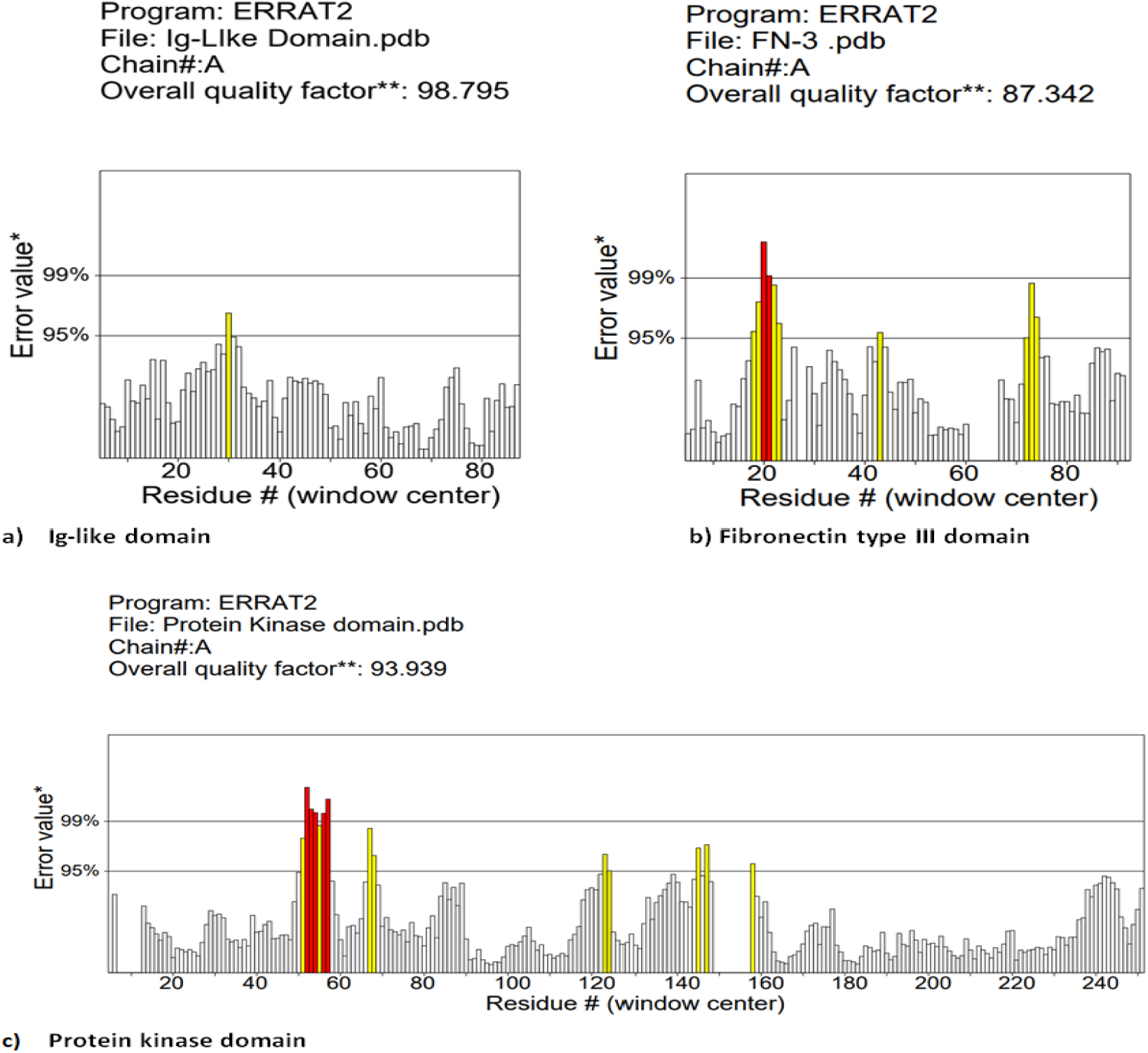
ERRAT 2 quality assessment of the studied domains. (a) Ig-like domain, (b) Fibronectin type III, and (c) protein kinase domain. These graphs illustrate residue wise error score derived from non-bonded atom-atom interactions. Yellow and red bars highlight regions exceeding the 95% and 99% error threshold, respectively. Most residues fall within the acceptable ranges supporting the structural validity of the models.

Structural validation through Ramachandran plot and ERRAT 2 confirms that the predicted structures for the studied domains exhibit reliable backbone geometry and acceptable non-bonded atomic interaction profiles.

### Hydrogen bonds Analysis

Hydrogen bonding significantly contributes to maintaining the β-sheet architecture, reinforcing loop structure, and supporting the global stability of protein structure [31]. In line with this, the studied Ig-like, Fn-III and Protein kinase domain showed significant hydrogen bonding pattern that help in predicting and comparing their stability for the estimation of degradation pattern for PMI estimation. Hydrogen bonding analyses were conducted using UCSF Chimera (v1.19), which automatically determines hydrogen bonds using predefined geometric parameters and considering the chemical nature of the involved atoms. By using refined algorithm, Chimera distinguishes true hydrogen bonds from steric clashes, facilitating precise identification of stabilized interactions across each titin domain [32]. The corresponding structural representations of each domain are presented in Figure 9, Figure 10, Figure 11.

**Figure 9.**
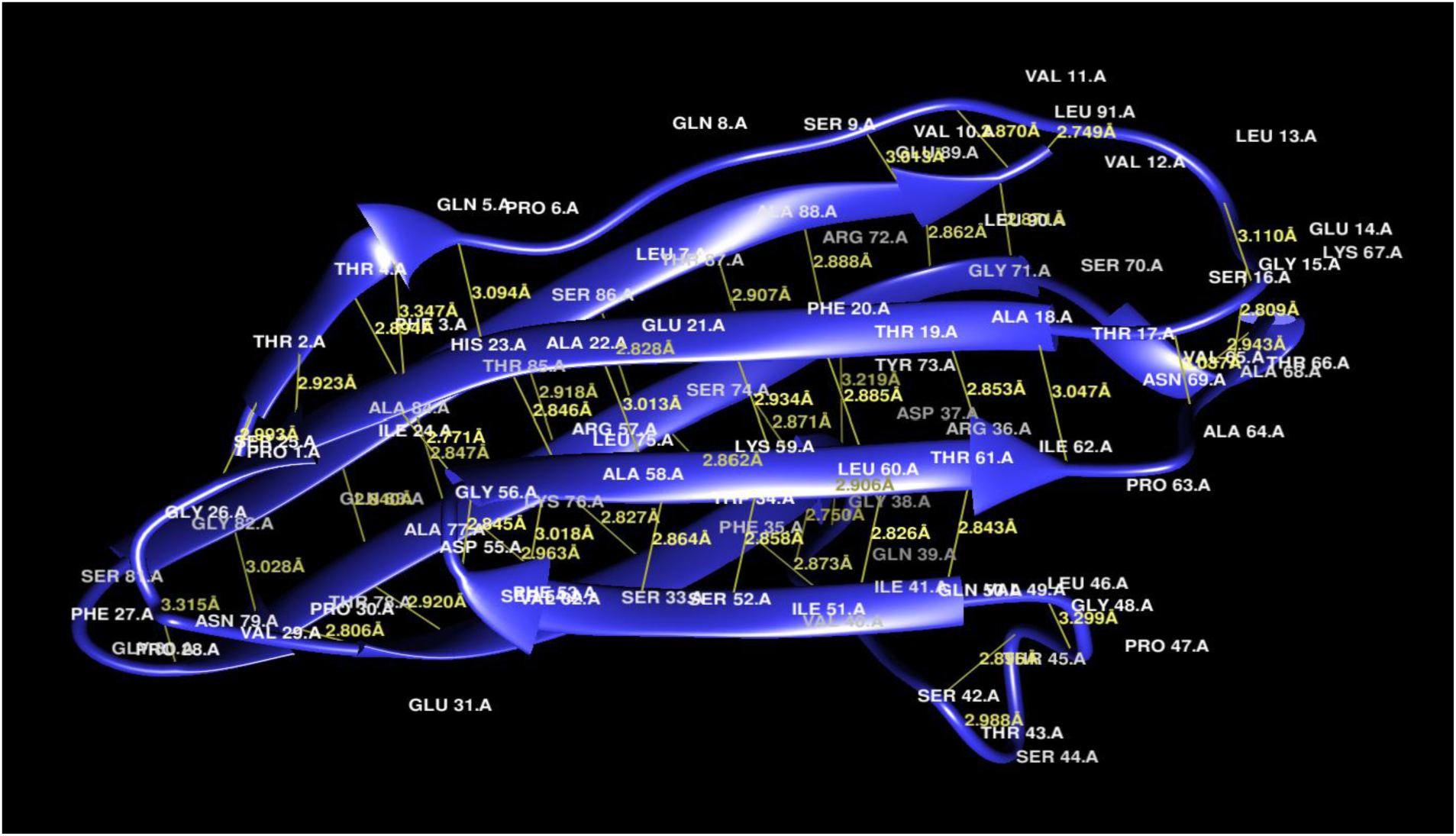
Visualization of hydrogen bond interactions within Ig-like domain of human titin protein. Structural representation of selected human titin domains. Figure 9 show ribbon model of the Ig-like, FN-III, and Protein kinase domains, respectively. Structured were visualized by using UCSF Chimera (v.1.19). Secondary structure elements are highlighted and hydrogen bonds are depicted as yellow dashes where applicable with distance in angstroms (Å).

**Figure 10.**
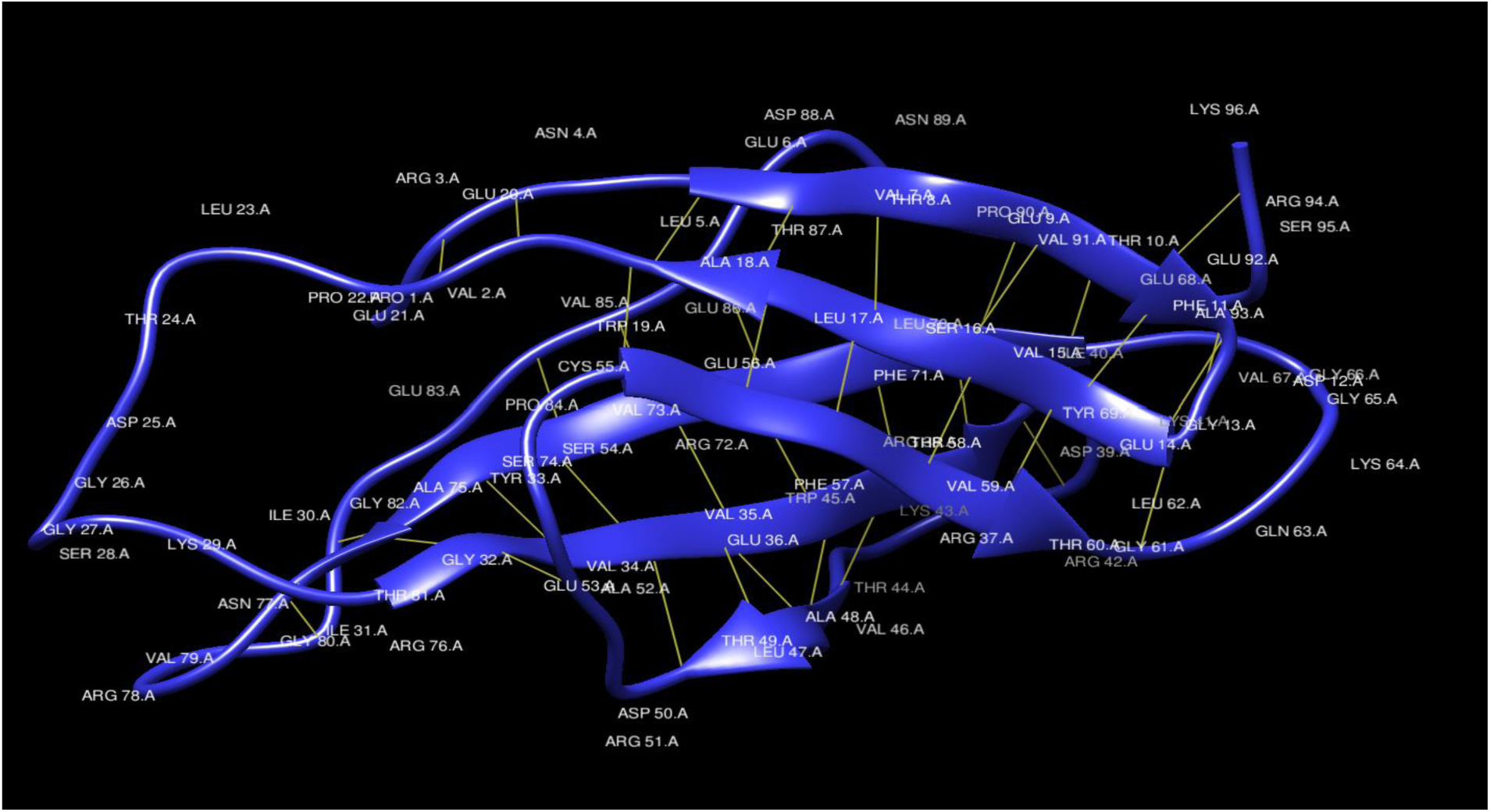
Visualization of hydrogen bond interactions within Fn-III domain of human titin protein. Structural representation of selected human titin domains. Figure 10 show ribbon model of the Ig-like, FN-III, and Protein kinase domains, respectively. Structured were visualized by using UCSF Chimera (v.1.19). Secondary structure elements are highlighted and hydrogen bonds are depicted as yellow dashes where applicable with distance in angstroms (Å).

**Figure 11.**
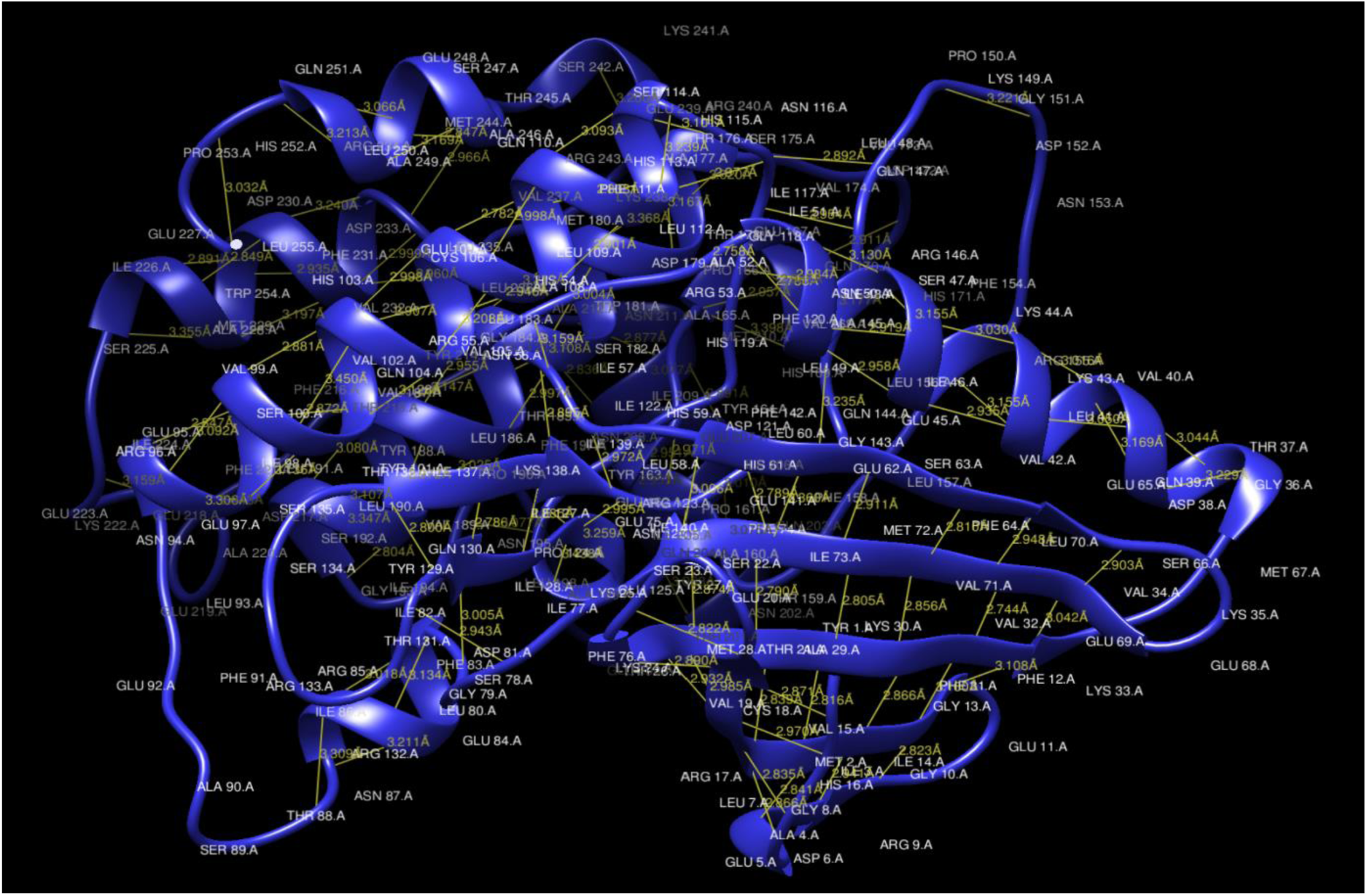
Visualization of hydrogen bond interactions within Protein kinase domain of human titin protein. Structural representation of selected human titin domains. Figure 11 show ribbon model of the Ig-like, FN-III, and Protein kinase domains, respectively. Structured were visualized by using UCSF Chimera (v.1.19). Secondary structure elements are highlighted and hydrogen bonds are depicted as yellow dashes where applicable with distance in angstroms (Å).

Hydrogen bonds were characterized by the most widely accepted geometric criteria established by McDonald and Thornton (1994). Within this framework, a hydrogen bond was considered present if the donor-acceptor distance is ≤3.5Å and the donor-H-acceptor angle is ≥ 120°, fulfilling basic stereochemical requirements [33]. On the basis of this study, hydrogen bonding interactions, a summary of total hydrogen bonds and Bonds having a distance less than ≤3.5Å of the studied domains are given in Table 4.

**Table 4.** Summary of the hydrogen bonding in the studied domains of human titin protein. This table shows the total numbers of hydrogen bonds observed and refining of the numbers (formation of subsets) according to the study of McDonald and Thornton indicating geometrically favorable interactions.

| **Protein Domain** | **Total H-Bonds** | **H-Bonds $\leq 3.5\text{\AA}$** | **Strong H-Bonds $\leq 3.2\text{\AA}$** | **Ratio of Strong to Total bonds** |
| --- | --- | --- | --- | --- |
| **Ig-like** | 67 | 66 | 57 | 0.851 |
| **Fibronectin Type III** | 67 | 66 | 58 | 0.866 |
| **Protein kinase** | 234 | 231 | 199 | 0.850 |

According to Table 4, this study concludes the order of stability, based on the ratio of strong to total hydrogen bonds is given as: Fibronectin type III is most stable, followed by Ig-like and Protein Kinase domain, respectively (FN- III > Ig-like > Protein Kinase).

## 3. Discussion

In silico approaches were employed, such as physicochemical properties, secondary structure, solvent accessibility, and 3D modeling. Ramachandran plot assessment, ERRAT scores, and hydrogen bond analysis were employed to determine the folding patterns and structural stability of the studied Titin domains. This study connects the structural stability and robustness of the selected domains to their postmortem persistence, addressing a critical gap in computationally driven forensic protein research. These studied features highlight which domains are more likely to remain stable and resistant to degradation processes as shown in Table 5, indicating their potential to use them as reliable molecular indicators for postmortem interval (PMI) estimation.

**Table 5.** Summary of computational analysis and predicted postmortem degradation behavior of selected titin domains. Physicochemical characteristics, solvent accessibility, structural features, and hydrogen bonding patterns were analyzed to infer relative postmortem proteolysis rates of Ig-like, FN-III, and protein kinase domains.

| **Sr#** | **Computational Properties** |  | **Ig-like Domain** | **Fn-III Domain** | **Protein kinase Domain** |
| --- | --- | --- | --- | --- | --- |
| **01)** | Physicochemical properties | Instability Index | 30.61 | 41.83 | 46.63 |
|  |  | Aliphatic Index | 81.43 | 78.02 | 95.57 |
|  |  | GRAVY Score | 0.004 | -0.537 | -0.163 |
|  |  | Estimated Half-life | >20 hours | >20 hours | 2.8 hours |
| **02)** | Solvent Accessibility | - | Mostly buried(0-2), slightly in (6-9) | Mostly buried, but some with $\geq 7$ | Mostly (0-3), and some with (4-6), and very less with >7 |
| **03)** | Structural Analysis | 2D-Structure | High α-helices and β-strands | Less α-helices and β-strand as compared | Moderate α-helices and β-strands |
| | | 3D-Structure | Very high β-<br>sheets with<br>minimum<br>loops and coils | High β-<br>sheets with<br>few loops<br>and coils | Moderate β-<br>sheets with<br>many loops and<br>coils |
| 04 | Hydrogen Bonding | - | 0.851 | 0.866 | 0.850 |
| 05) | Interpretation | - | Slow<br>proteolysis<br>after death;<br>expected to<br>degrade at last | Moderate<br>proteolysis<br>after death;<br>expected to<br>degrade<br>gradually | Rapid<br>proteolysis after<br>death; expected<br>to degrade early |

By providing a reliable computational reference for domain stability, this study bridges a major methodological gap in forensic proteomics, where in silico evaluations are still limited. Researchers used this study as a reference for aligning wet-lab postmortem for more accurate interpretation and strengthening protein-based PMI analyses.

Previous studies have analyzed proteins as a forensic marker [34,35] but none of them focused on comprehensive domain-level computational analysis. Existing proteomics studies related to PMI approaches don’t focus on titin domains, so this study proposes titin domains as a reliable candidate for PMI estimation. To the best of current studies, this is the first in silico structural analysis of titin domains designed for PMI estimation, providing a novel framework for further experimental studies and development of titin-based forensic assays.

### Comparison of Previous Postmortem Proteins studies

In order to support the in-silico findings of Titin domains as a potential PMI marker, selected literature examining postmortem muscle protein degradation relevant to PMI estimation is compiled in Table 6.

**Table 6.**
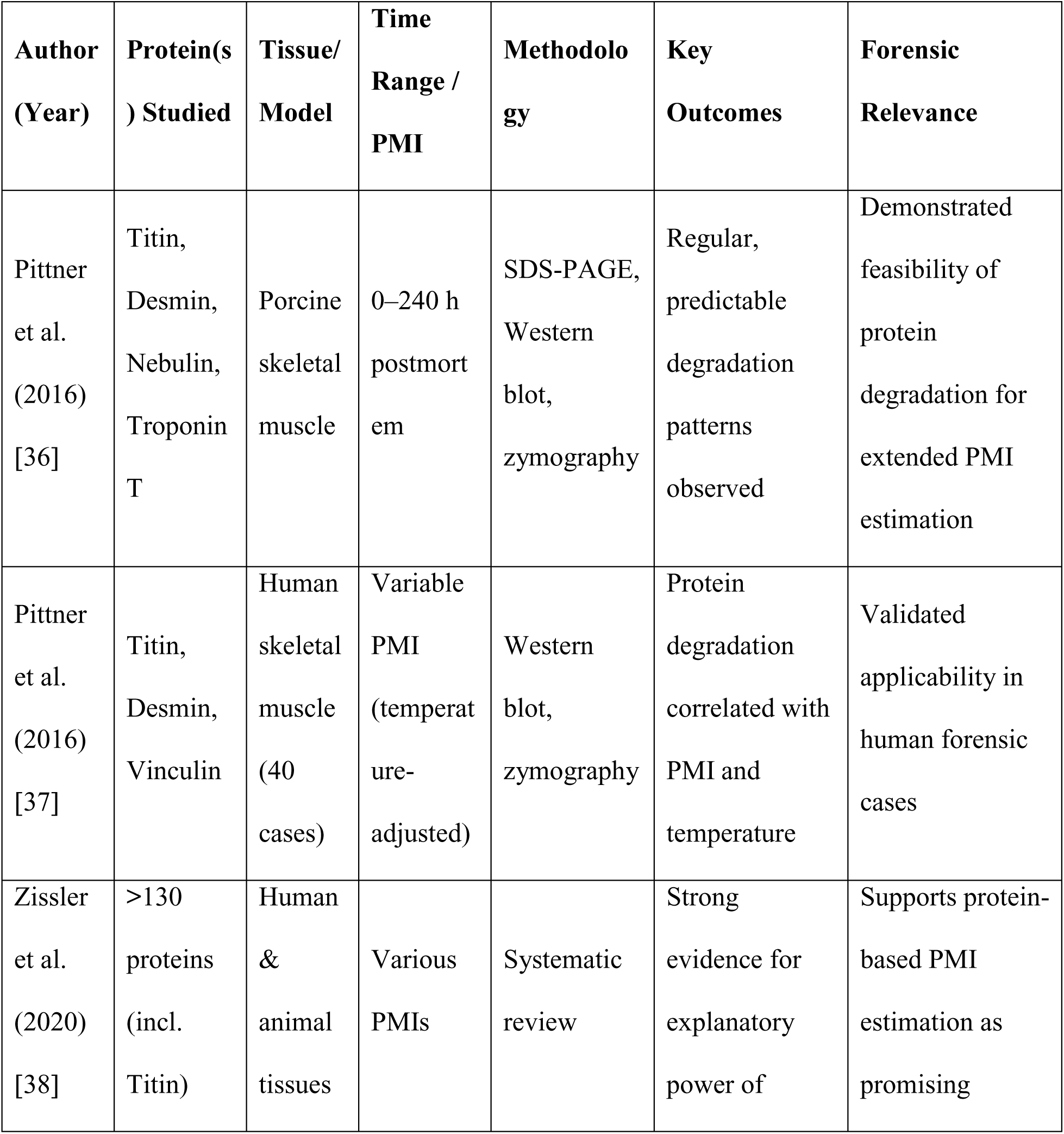
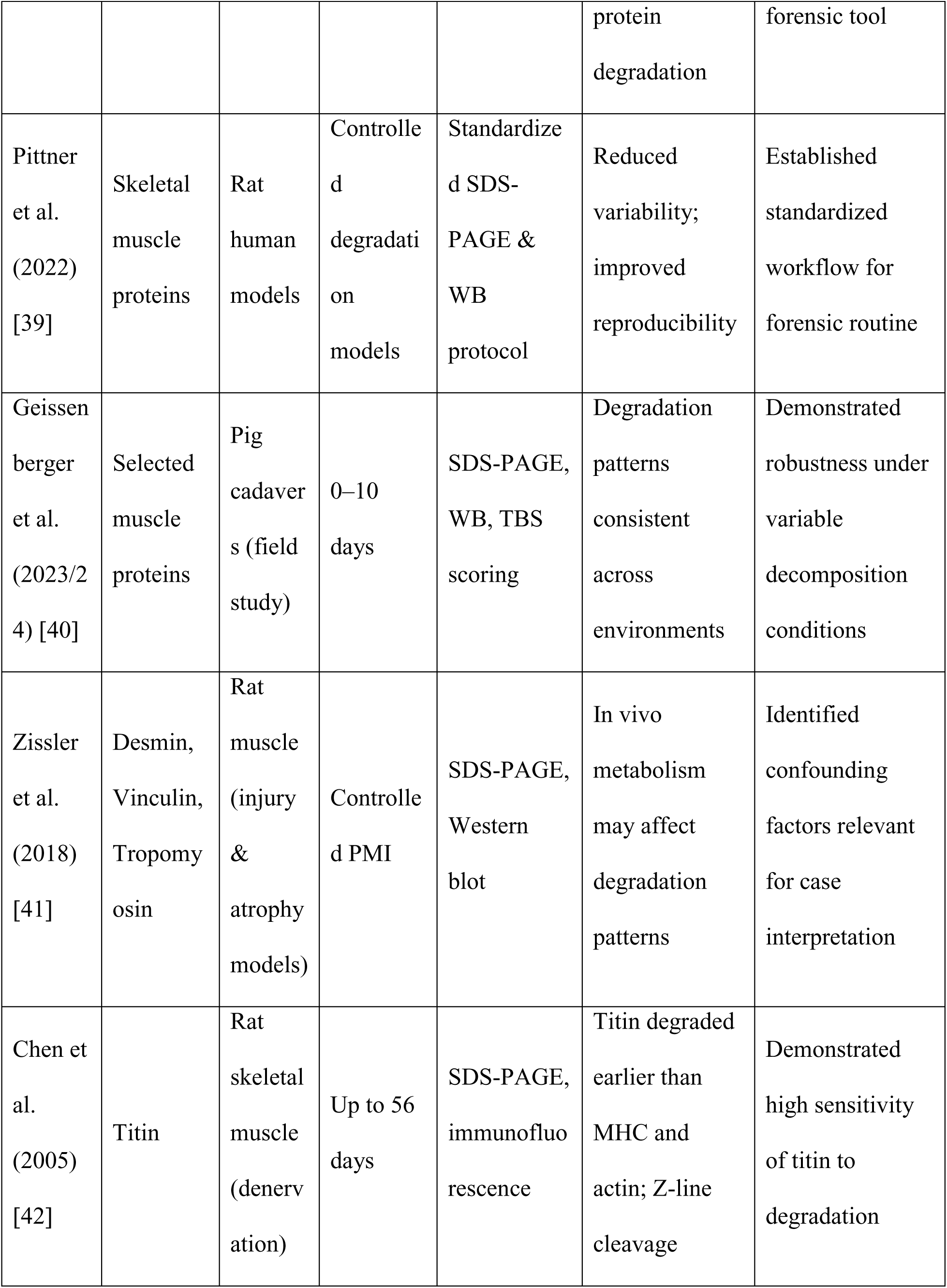
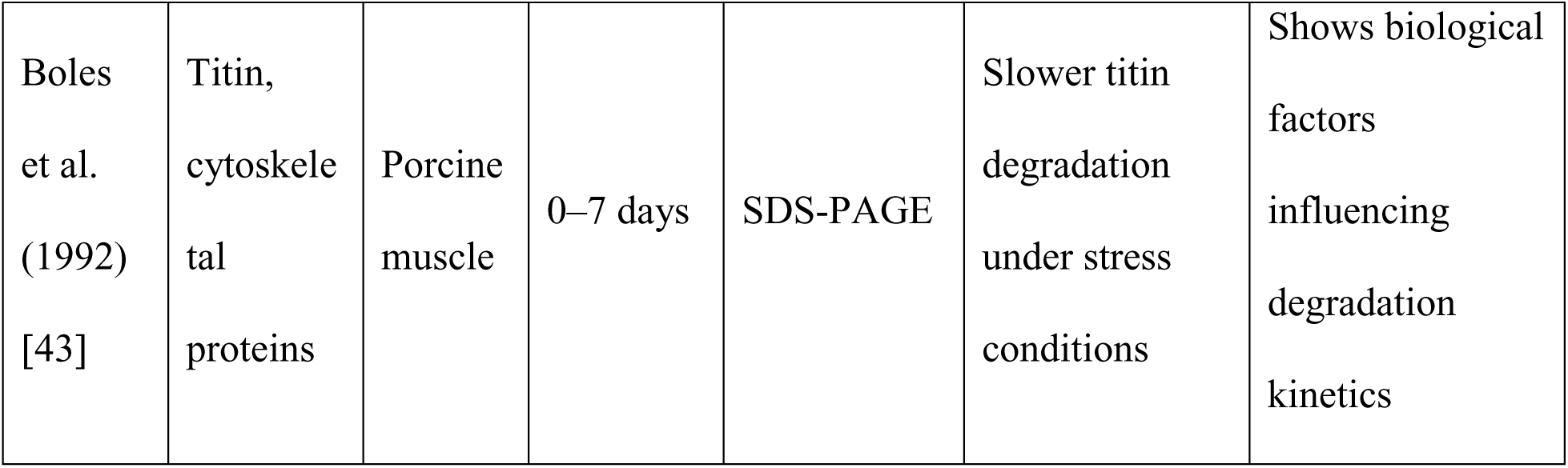
Summary of selected literature reporting postmortem degradation pattern of skeletal muscle proteins relevant to PMI estimation. PMI, postmortem interval; SDS-PAGE, sodium dodecyl sulfate–polyacrylamide gel electrophoresis; WB, Western blot; TBS, total body score; MHC, myosin heavy chain. Time ranges refer to postmortem intervals investigated under controlled or field conditions. Listed studies include both animal and human models commonly used to evaluate skeletal muscle protein degradation patterns relevant to forensic PMI estimation.

Previous experimental investigations have consistently demonstrated that large skeletal muscle proteins undergo time-dependent postmortem degradation, providing valuable information for (PMI) estimation. In particular, titin has been repeatedly identified as a highly sensitive and early-degrading structural protein across both animal models and human forensic cases. Studies employing SDS-PAGE, Western blotting, and zymographic approaches have shown reproducible degradation patterns of titin that correlate with PMI, temperature, and decomposition conditions, supporting its forensic relevance over extended postmortem periods Table 6. Collectively, these findings establish a strong biological and forensic foundation for considering titin as a promising PMI-related marker. Building upon this experimental evidence, the present in-silico study focuses on domain-level characterization of titin to explore structural and physicochemical features that may underlie its postmortem degradation behavior and further support its potential application in forensic PMI estimation.

## 4. Conclusion

This research provides an in-depth computational analysis of human Titin domains to assess their stability as a molecular marker for postmortem interval estimation (PMI) estimation. This analysis identifies distinct stability profiles across the domains that Ig-like is most stable followed by Fn-III and Protein kinase domain which govern their rate of post-mortem degradation. By addressing the gap in domain-specific titin modeling in prior forensic literature, this study introduces a foundation for using titin in PMI-oriented biomarker development.

## 5. List of abbreviations

PMI: Postmortem interval estimation
Ig-like: Immunoglobulin like
Fn- III: Fibronectin Type III
PDB: Protein data bank
PSIPRED: Position-Specific Iterated Prediction of Secondary Structure
SWISS-model: Swiss Institute of Bioinformatics Protein Structure Homology-Modelling Server
ExPASY: Expert Protein Analysis System
GRAVY: Grand Average of Hydropathy
QMEAN: Qualitative Model Energy Analysis
GMQE: Global Model Quality Estimation
ERRAT: Error Assessment of Protein Models
UCSF Chimera: University of California, San Francisco Chimera

## 6. Declarations

### Ethics approval

Ethical approvals are not applicable because it is computational study not the wet lab. This study is pure in silico and does not involve human participation, human tissues, human data, and any kind of living organism.

### Consent for publication

Not applicable

### Funding

No external funding was received for this study.

## Notes

### Competing Interest Statement

The authors have declared no competing interest.

